# miR169 and *PmRGL2* synergistically regulate NF-Y complex to active dormancy release in Japanese apricot (*Prunus mume* Sieb. et Zucc.)

**DOI:** 10.1101/2020.02.24.963355

**Authors:** Gao Jie, Ni Xiaopeng, Li Hantao, Faisal Hayat Make, Shi Ting, Gao Zhihong

## Abstract

Insufficient chilling requirements affect the floral bud quality and fruit yield in fruit crop production. Endodormancy is a process to meet the chilling requirement. To understand the mechanism of dormancy release in woody plants, we compared the miRNA database during the transition stage from endodormancy to dormancy release in Japanese apricot and found that the miR169 family showed significant differentially up-regulated expression during dormancy and down-regulated during dormancy release periods. The 5’ RACE assay and RT-qPCR validated its target gene NUCLEAR FACTOR-Y subunit A (NF-YA) exhibited the opposite expression pattern. Further study showed that exogenous GA_4_ could inhibit the expression of *PmRGL2* and promote the expression of *NF-Y*. Moreover, the interaction between NF-Y family and GA inhibitor *PmRGL2* was verified by yeast-two-hybrid system and Bimolecular fluorescence complementarity (BiFC) experiment. These results suggested that synergistic regulation of NF-Y and *PmRGL2* complex to active dormancy release induced by GA_4._ These will help to elucidate the functional and regulatory roles of miR169 and its target gene of the seasonal bud dormancy induced by GA_4_ in Japanese apricot woody plants and to provide new sights for the discovery of dormancy release mechanism.

## Introduction

In winter, the perennial woody trees, including deciduous fruit trees, have to face the most severe threats like low temperature and short photoperiod intemperate zones. Flower bud dormancy is a bet-hedging strategy to prevent injury in inappropriate circumstances and then synchronize of perennial growth (Zhang et al., 2018). The release of seasonal dormancy requires a certain amount of chilling requirements (Benmoussa et al., 2017;Prudencio et al., 2018) or dormancy breaker (Posner et al., 2018), such as gibberellin (Rinne et al., 2011; Saure, 2011). Furthermore, bud dormancy influence the flowering time and quality of flower buds, which will affect the yield and quality of fruits (Campoy et al., 2011; Steven et al., 2014). Lang et al. (1987) had divided the pattern of dormancy into three periods, paradormancy, endodormancy and ecodormancy. The low temperature in autumn causes deciduous plants to enter the phase of paradormancy. After the leaf falling, these buds shift into the endodormancy state. At this period, the dormancy cannot be released even under favourable conditions. When plants entered into ecodormancy, the dormancy release can be induced by external environmental factors (Wisniewski et al., 1996; Horvath et al., 2003). Japanese apricot (*Prunus mume* Siebold et Zucc.) originated in China and determined as earliest flowering deciduous fruit trees with the diverse chilling requirement from 26.3 to 75.7CP (Zhuang et al., 2016). So it is an excellent plant material for studying the mechanism of seasonal dormancy in fruit trees.

MicroRNAs are the short non-coding RNAs (21–24 nucleotides) (Axtell and Meyers, 2018) that regulate the expression of target genes through complementary sequence matching. In plants, miRNAs regulate the expression of various genes in cluding transcription factors (Jung et al., 2014; Hernandez et al., 2017), stress-responsive proteins (Jagadeeswaran, G et al. 2009; Sun, X. et al. 2015), and other proteins, which are involved in the processes of growth and development (Jonesrhoades et al., 2006; Curaba et al., 2013; Chen et al., 2019), disease resistance (Feng et al., 2012; Yu et al., 2017; Jiang et al., 2018), abiotic stress response (Zhan et al., 2012; Zheng et al., 2015; Singh et al., 2017) and physiology (Gao et al., 2012a; Ma et al., 2019). Although the regulation of genes by miRNA is indirect, it is believed that miRNAs play essential roles in plant development.

Multiple studies have shown that miR169s targets of the NF-YA (CBF-B or HAP2) transcription factors (Hanna et al., 2010; Francesco et al., 2011; Meng et al., 2011; Gyula et al., 2018). One of the subunits of the NF-Y family, which together with two other subunits NF-YA and NF-YC, forms a heterotrimeric complex, binds to the CCAAT box of downstream genes to activate or inhibit their expression (Antonella et al., 1995; Liu et al., 2016; Martyn et al., 2016). All of which are required for DNA binding (Mantovani 1999). In recent years, the function of miR169 has been reported in different species. Meng et al. (2011), which revealed that miR169a was involved in nitrogen starvation of *Arabidopsis thaliana*, and transgenic miR169a contained less nitrogen than the wild type. It was reported that miR169 gene was regulated by the development of symbiotic nodules. Specific miR169 isoforms and NF-YA2 target control root structures in *Arabidopsis thaliana* were proposed (Céline et al., 2014). Luan et al. (2015)also found that miR169s and its target NF-YA were involved in response to abiotic stress in maize leaves. The similar resultswere found in soybean, where the target gene *GmNF-YA3* of miR169 was a positive regulator of plant tolerance to drought stress (Ni et al., 2013). Some reportssuggest that miR169 targets NF-YA and participates in the bud dormancy (Rewati et al., 2013; Ding et al., 2016).

GA_4_ is synthesized from the respective precursors, GA_12_, after a few oxidations by GA20-oxidase (GA20ox) and final oxidation by GA3-oxidase (GA3ox) (Sven et al., 2006). Rinne et al. (2011) consideredthe function of GA_3_ and GA_4_ in dormancy is distinguished, and only GA_4_ can better replace part of the chilling requirement. In the previous studies, our group has proved that GA_4_, as a good dormancy breaker, can replace a part of chilling requirements and cause an early release of dormancy in Japanese apricot (Zhuang et al., 2013). DELLA (Asp-Glu-Leu-Leu-Ala) is a crucial negative regulatory element of the GA signal transduction pathway, which is a subfamily of the GRAS gene family (Salanenka et al., 2018; Nimisha et al., 2013). It can regulate the expression of some transcription factors related to plant growth and thus inhibit plant growth (Peng et al. 1997; Acheampong et al., 2017;Chen et al., 2017;Zhang et al., 2018). Recent studies have shown thata DELLA–EDS1-mediated feedback regulatory loop by which plants maintain the subtle balance between growth and defense to avoid excessive growth or defense in response to constant biotrophic pathogen attack (Y, Li et al., 2019). In Arabidopsis, the DELLA family mainly consists of five highly homologous protein inhibitors, namely GA-INSENSITIVE (GAI), REPRESSOR OF ga1-3 (RGA), RGA-LIKE 1 (RGL1), RGL2 and RGL3(Suzuki et al., 2009). The binding of *RGL2* protein and DOF6 protein (DNA BINDING1 ZINC FINGER6) can activate the expression of GATA, thus forcing seed dormancy (Ravindran et al., 2017). Dmitriev et al. found that *RGL2* is not only a negative regulator of GA-mediated seed germination in *Arabidopsis thaliana* but also related to GA-induced flower development (Dmitriev et al., 2019).

In Japanese apricot, *PmDAMs* gene has been shown to play an essential role in dormancy (Ryuta et al., 2011; Yamane and Tao, 2015; Zhao et al., 2018). *PmCBF* can interact with *PmDAM6* in dormancy induction (Zhao et al., 2018). Transcriptome profiles reveal the critical roles of hormone and sugar in the bud dormancy (Zhang et al., 2018). In order to study the molecular mechanism of dormancy, our group investigated the chilling requirement and heat requirement of 69 different Japanese apricot varieties, determined the requirements of cold storage capacity during dormancy, and used proteomics, transcriptomics, and metabolomics to study each dormancy period (Zhuang et al., 2012). *PmRGL2* inhibits dormancy release through the GA signaling pathway, and overexpression of *PmRGL2* showed slow growth and a late germination phenotype compared to the control (Lv et al., 2018). However, the mechanism of dormant molecular regulation is not completely clear. It is necessary to study the mechanism of dormancy in fruit trees. In this study, we found that small RNA played a vital role in the dormancy shift. Among them, miR169 family target to *PmNF-YA3* gene, which has noticeable changes in dormancy and dormancy release. Direct cleavage of *PmNF-YA3* mRNA by miR169 was confirmed by 5 ‘RACE assays. Further, validation revealed that*PmNF-YA3, PmNF-YB3, PmNF-YC1* interact with *PmRGL2* in vivo, indicating its essential role in the GA signal transduction during dormancy.

In order to further understand the molecular mechanism of dormancy in Japanese apricot, we used high-throughput sequencing of small RNAs about the four critical stages of Japanese apricot flower buds (Para, Endo, Eco, and Dr) to discover the regulatory roles of small RNA presumably involved in the dormancy.

## Results

### miRNA high-throughput sequencing and characterization of Pmu-miR169 family members

In this study, high-throughput miRNA sequencing was performed on the flower buds of Para, Endo, Eco, and DR periods to dig out the miRNAs involved in dormancy. The significant differentially expressed miRNAs and their targeted transcription factors in dormancy induction and dormancy release transition stages are shown in Table S2. It is interesting that the expression of miR169 family was significantly up-regulated in dormancy induction and down-regulated in dormancy release shown in the Table 1, which suggested that it might play a crucial role in the dormancy shift.

**Table 1:**
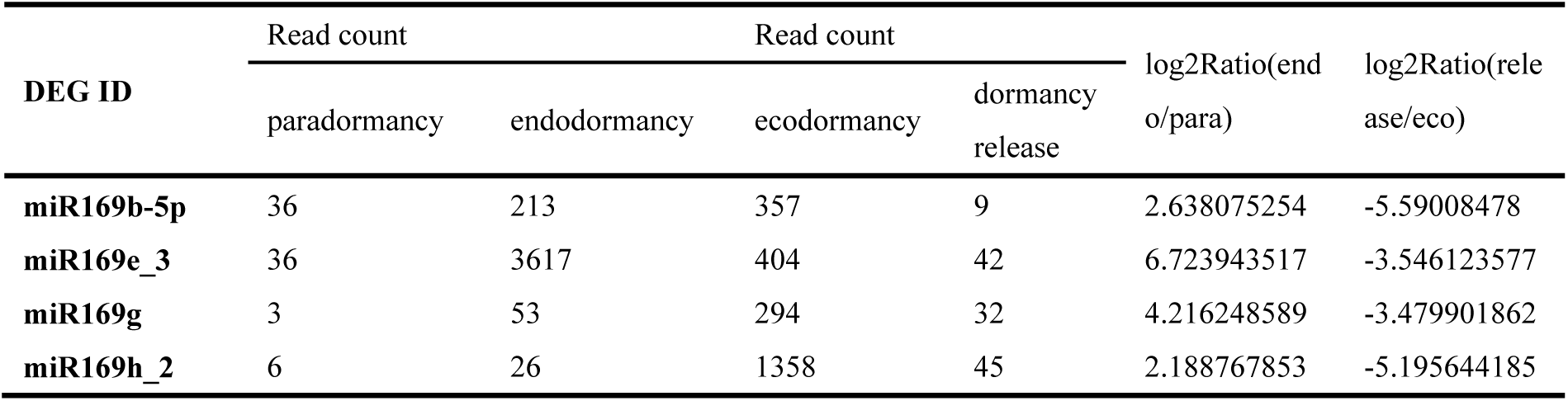
The expression pattern of miR169s during the four stages of dormancy in Japanese apricot.

Based on the analysis of the miRNAs database, eleven Pmu-miR169 genes were located on chromosomes 4 in Japanese apricot. A phylogenetic tree based on the full-length premiR169 sequences showed that the Pmu-miR169c-3p_2 and Pmu-miR169d-5p_1 precursors, which generated the same mature sequence, shared high homology (Fig.1). Similarly, high homology was exhibited between the Pmu-miR169e_3 and Pmu-miR169w precursors. However, the Pmu-miR169g, and Pmu-miR169h_2 precursors, and the Pmu-miR169k and Pmu-miR169v_1 precursors, Pmu-miR169f_1 with Pmu-miR169c-3p_2 and Pmu-miR169d-5p_1, showed low similarity within each group although the members of each group generated the same mature Pmu-miR169s, suggesting that they might regulate the same target genes at different development stages or in different tissues.

**Figure 1:**
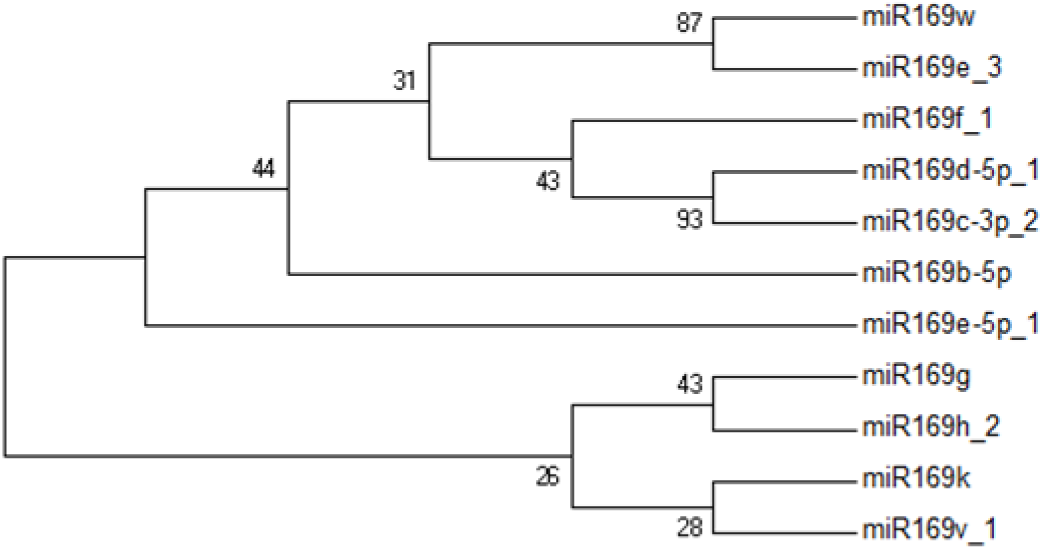
A phylogenetic tree of 11 miR169s family members from Japanese apricot.

### 5’RACE experiment validated the targets of miRNA and related gene expression levels

#### 1. Validation of Pmu-miR169 targets using 5’ RACE assay and expression of related genes

miRNAs participate in various biological pathways via silencing of gene expression in a targeted manner (Jean-Philippe et al., 2006). Therefore, the identification of the miR169 target is essential to understand its role in response to dormancy. Thus, the coding DNA sequence of Japanese apricot (CDs) was downloaded from NCBI (https://www.ncbi.nlm.nih.gov/) and by psRobot, TAPIR\reference (TAPIR_ref) and Target Finder to identify the potential targets of miR169. The results indicated that NF-YA is a candidate target of miR169. A potentially more reliable method for predicting miRNA and its targets, the results of miRNA and mRNA transcriptome sequencing were combined to screen useful targets according to the similarity of expression level changes (Dmitriev et al. 2013) (Fig.3). Among them, one *NF-YA1*, one *NF-YA10*, and three *NF-YA3* were predicted to be the target of miR169s, and the expression trends were contrary to it.

**Figure 2:**
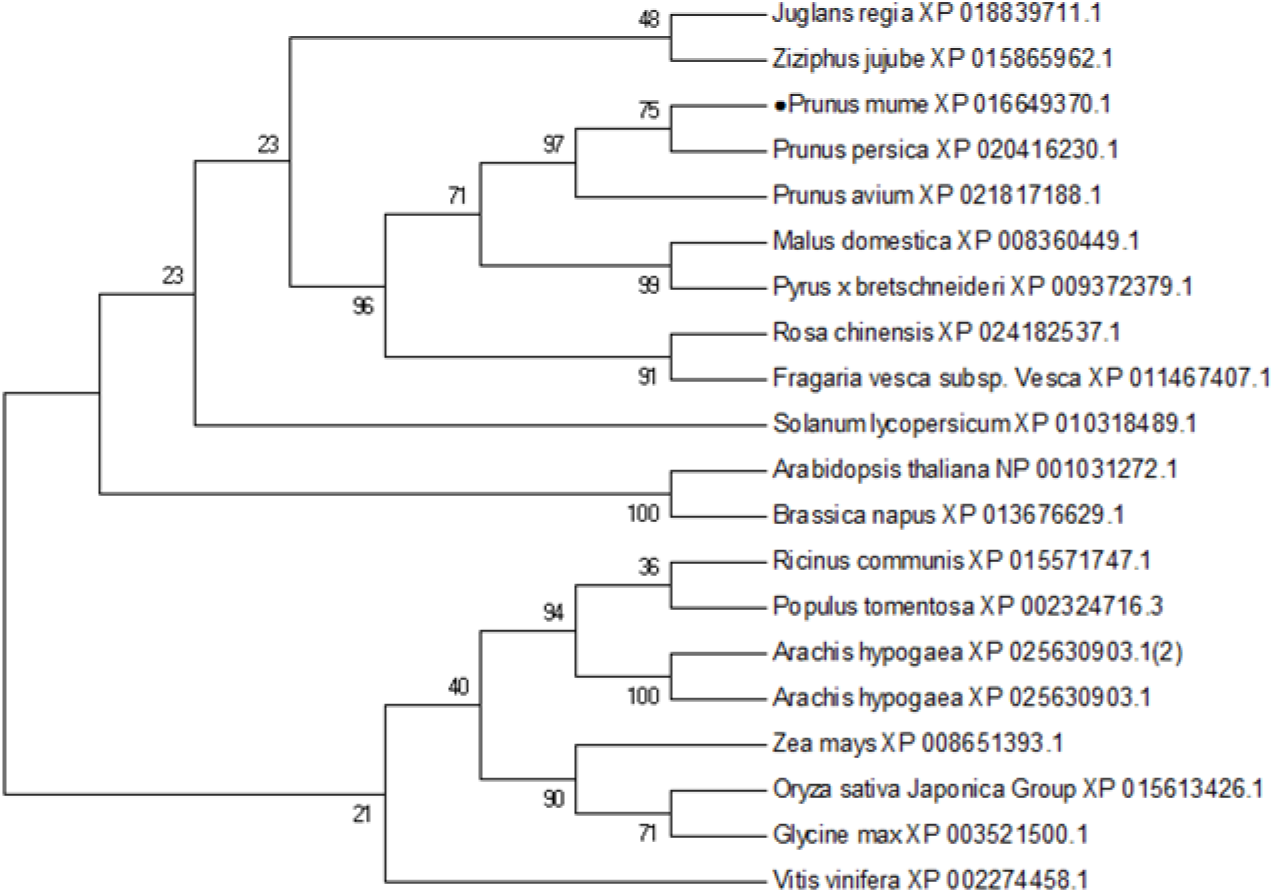
A phylogenetic tree of *PmNF-YA3* from Japanese apricot and NF-YA3 proteins from other plant species.

**Figure 3:**
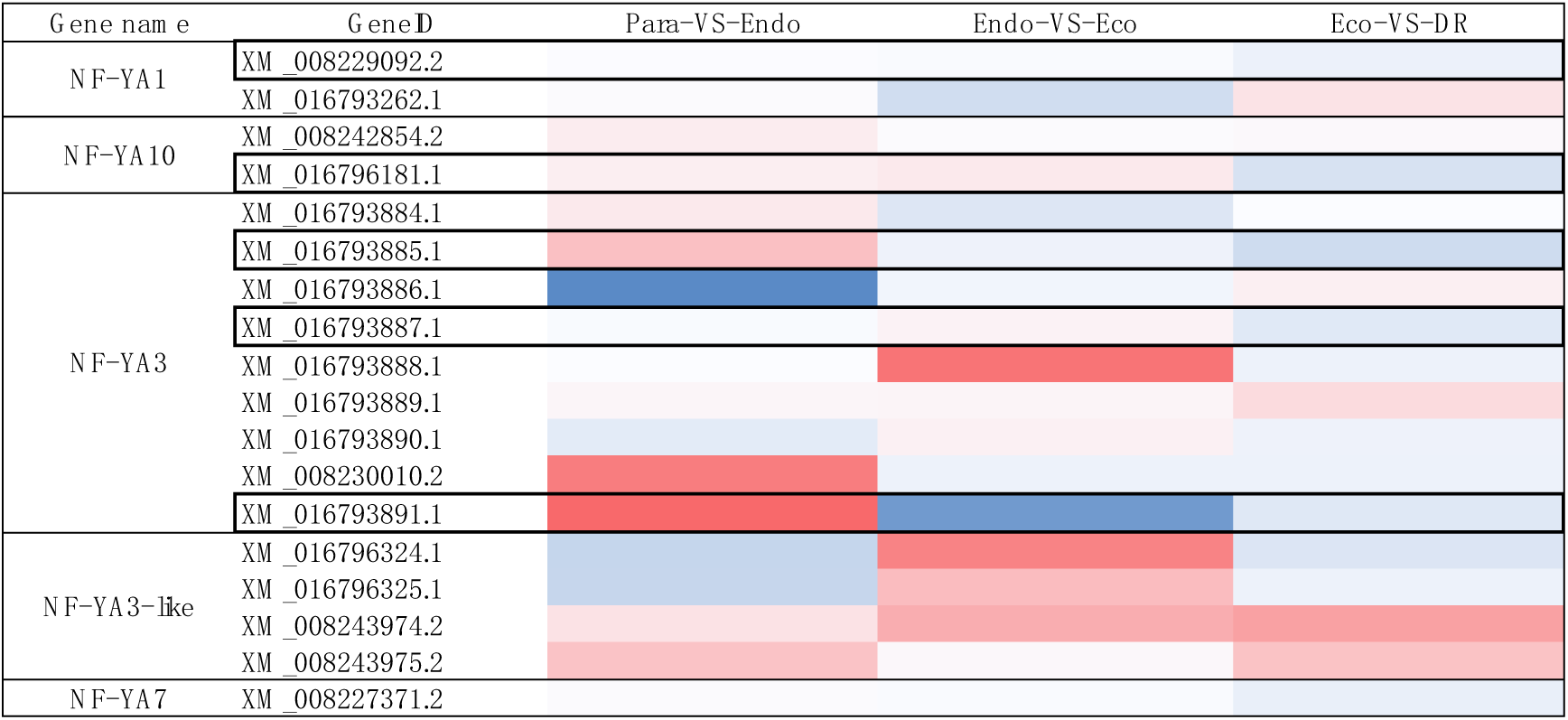
Cluster analysis of gene expression based on log ratio RPKM data. The cluster display expression patterns for a subset of DEGs in three comparisonsof NF-YA genes. (Para vs Endo, Endo vs Eco and Eco vs DR). Each column represents an experimental condition, and each row represents a gene. Red means up-regulated and blue means down-regulated.

In order to verify the accuracy of the computational prediction, a 5′ RNA ligase-mediated rapid amplification of cDNA ends (RLM-RACE) analysis was performed. The results confirmed that *PmNF-YA3* is the target gene of miR169. The identification of the accurate cleavage site indicates that the cleavage occurs between the 9^th^ and 10^th^ nucleotides of the target site (Fig. 4). A comparison of *PmNF-YA3* protein sequences with those of 19 other species showed that *NF-YA3* was ubiquitous in all species and closely related to Rosaceae (Fig. 2).

**Figure 4:**
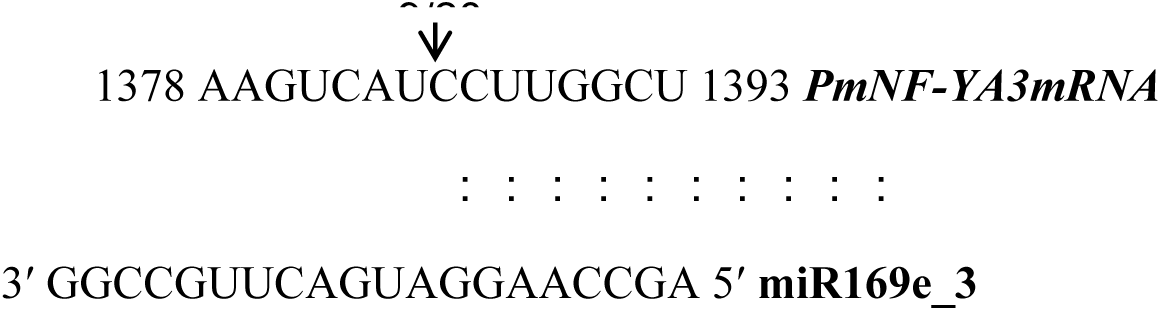
Identification of the predicted miR169 target site as determined with 5′ RLM-RACE. Notes: The position of 5′ RLM-RACE products are indicated with an arrow. The numbers above the sequence indicate the detected cleavage site in independent clones.

#### 2. Expression levels of miR169s and the related genes changed during the four periods of dormancy in Japanese apricot

Sequencing results revealed that the expression level of miR169s generally increased during dormancy period while decreased during the dormancy release period, indicating that miR169s played a positive regulatory role in dormancy. miR169s in four periods of dormancy were verified by RT-qPCR, which was consistent with the sequencing results (Fig. 5). At the same time, we tested the expression of NF-Y family members of flower buds during various stages of dormancy. The target gene *PmNF-YA3* of miR169s and its heterogeneous *PmNF-YB3* displayed the opposite expression trend to that of miR169s. However, *PmNF-YC1* showed different expression patterns from the other two subgroups, and it indicates that three subfamilies might play different roles in dormancy. This, we found that *PmNF-YC1* and *PmRGL2* showed a similar expression pattern (Fig. 6), and there may be a close relationship between them.

**Figure 5:**
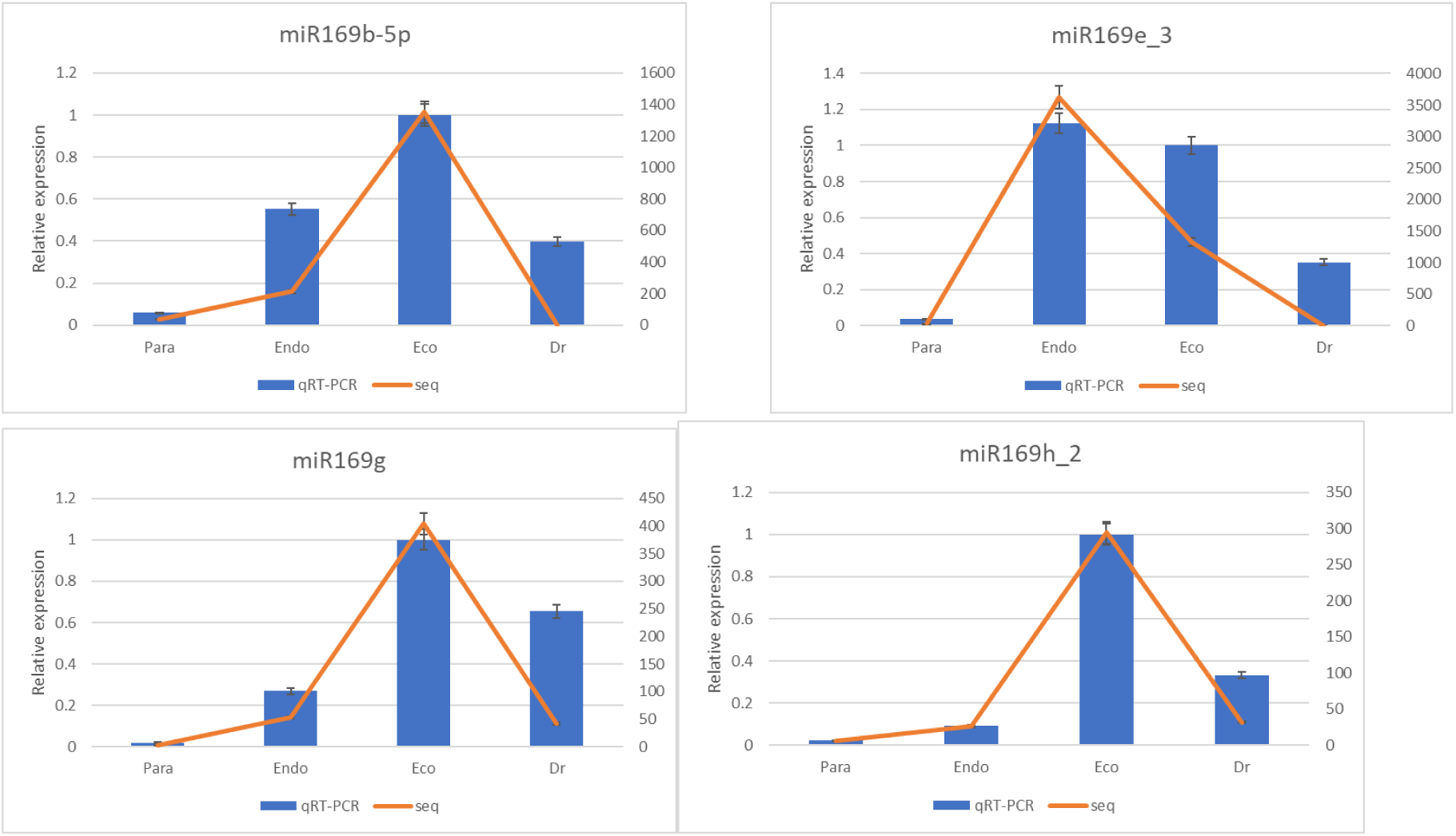
Expression changes miR169s in floral buds of Japanese apricot during the four stages of dormancy are determined by RT-qPCR. The y-axis shows the expression levels: the left side shows the relative expression levelas determined by RT-qPCR, whereas the right shows the tag number by DGE. Data represent the mean of three biological replicates where each biological replicate consisted of three technical replicates.

**Figure 6:**
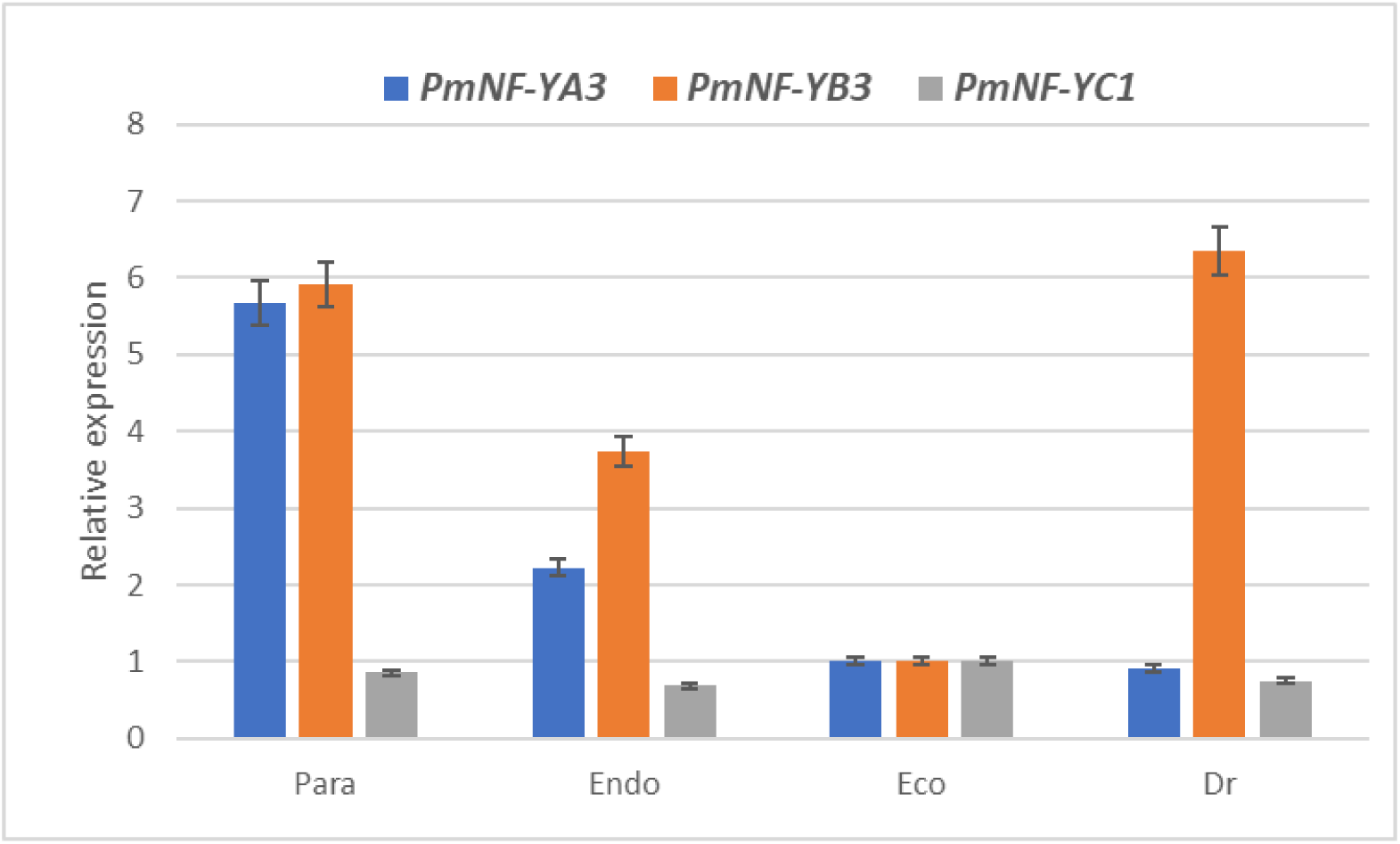
Expression changes NF-Y family members in floral buds of Japanese apricot during the four stages of dormancy are determined by RT-qPCR. Data represent the mean of three biological replicates where each biological replicate consisted of three technical replicates.

### Exogenous GA_4_ increased NF-Y expression

miR169 targets *NF-YA* gene by acting during dormancy, suggesting that the NF-Y family may play a role in dormancy. In order to confirm this conjecture, we broke the sticks of Japanese apricot in the ecodormancy period, treated them with the GA_4,_ and collected flower buds at 0, 3, 5, 8, and 10 days to detect the expression level of related genes. After ten days of treatment, the germination rate of the experimental group had reached 50%, where the dormancy was considered to be released, while the control group was fell far short. As shown, GA_4_ treatment significantly reduced the expression of *PmRGL2* compared with the control, while the expression of *PmNF-YA3, PmNF-YB3*, and *PmNF-YC1* showed up-regulation (Fig. 7). However, such results were more obvious in the early stage of treatment, and the expression level of *PmNF-YB3* and *PmNF-YC1* was lower than that of the control group after 5d treatment, which could be related to other molecular mechanisms.

**Figure 7:**
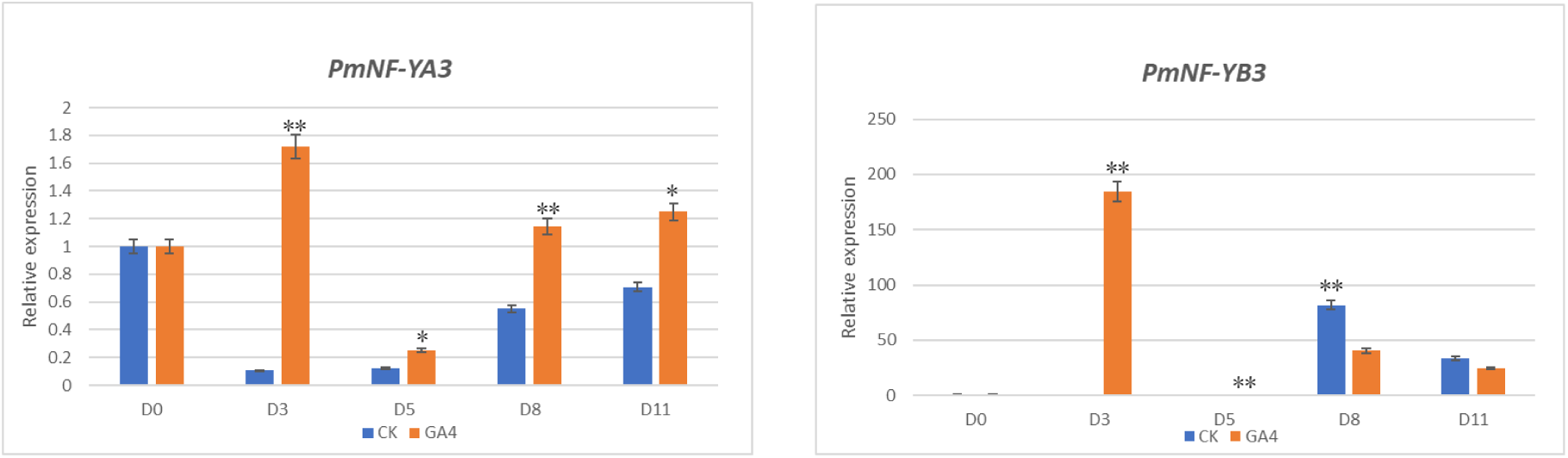
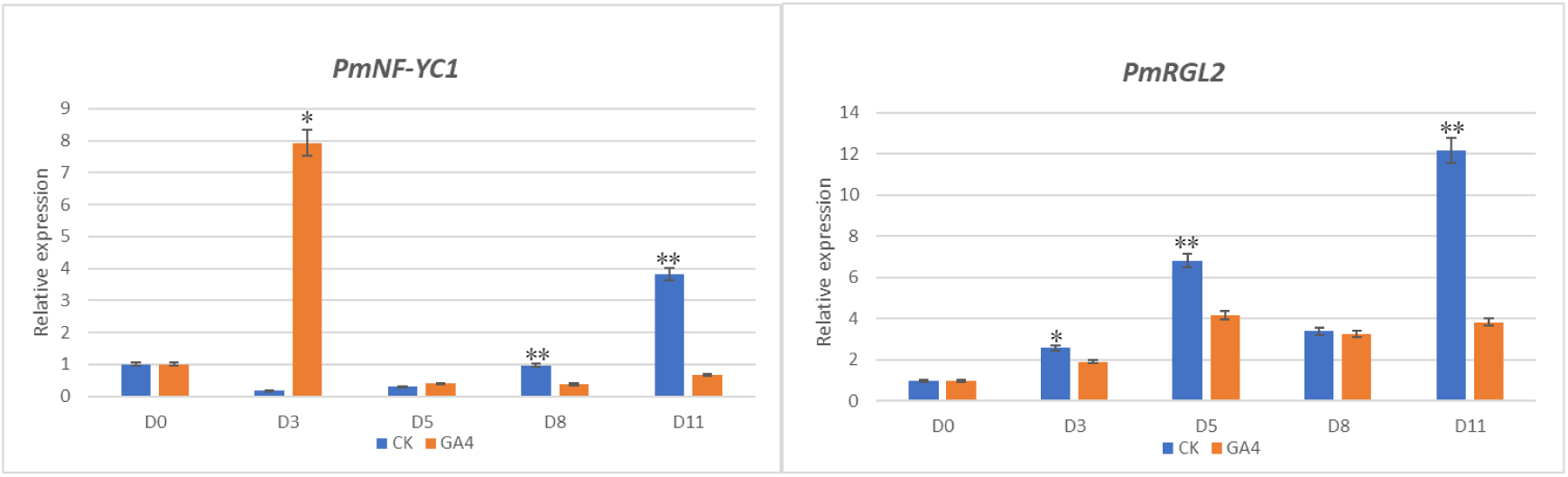
Changes in the expression of GA_4_ treated genes. Expression changes NF-Y family members and *PmRGL2* in floral buds of GA_4_ treated are determined by RT-qPCR. Significant differences between the experimental group and control group indicated by asterisks (*P≤0.05; **P≤0.01). (Duncan’s multiple range test). Data represent the mean of three biological replicates where each biological replicate consisted of three technical replicates.

### The NF-Y family interacts with *PmRGL2*

GA_4_ can promote the expression of the NF-Y family, so whether it is also involved in the release of GA_4_ induced dormancy release. In order to further understand the significance of NF-Y in dormancy release. The full-length CDs of *PmNF-YA3, PmNF-YB3*, and *PmNF-YC1* were amplified by PCR with gene-specific primers and inserted into pGADT7 and pGBKT7 vector, respectively. Transformed yeast with lab-stored pGBKT7-*PmRGL2*. Gene and no-load co-transformation grew on DDO but did not grow on QDO and QDO/A/X, like negative control, indicating that the gene could not be self-activated. Gene co-transformation can grow on both DDO and QDO, and appears blue on QDO/A/X, indicating *PmRGL2* can interact with *PmNF-YA3, PmNF-YB3*, and *PmNF-YC1*. Furthermore, *PmNF-YA3, PmNF-YB3* and *PmNF-YC1* can interact with each other (Fig.8). BiFC analysis verified the interactions among NF-Y subunits and between NF-Y subunits and *PmRGL2* in onion epidermal cells (Fig. 9). The results showed that PmRGL2 protein interacted with NF-Y protein subunits and NF-Y protein subunits interacted with each other in the nucleus of Japanese Apricot.

**Figure 8:**
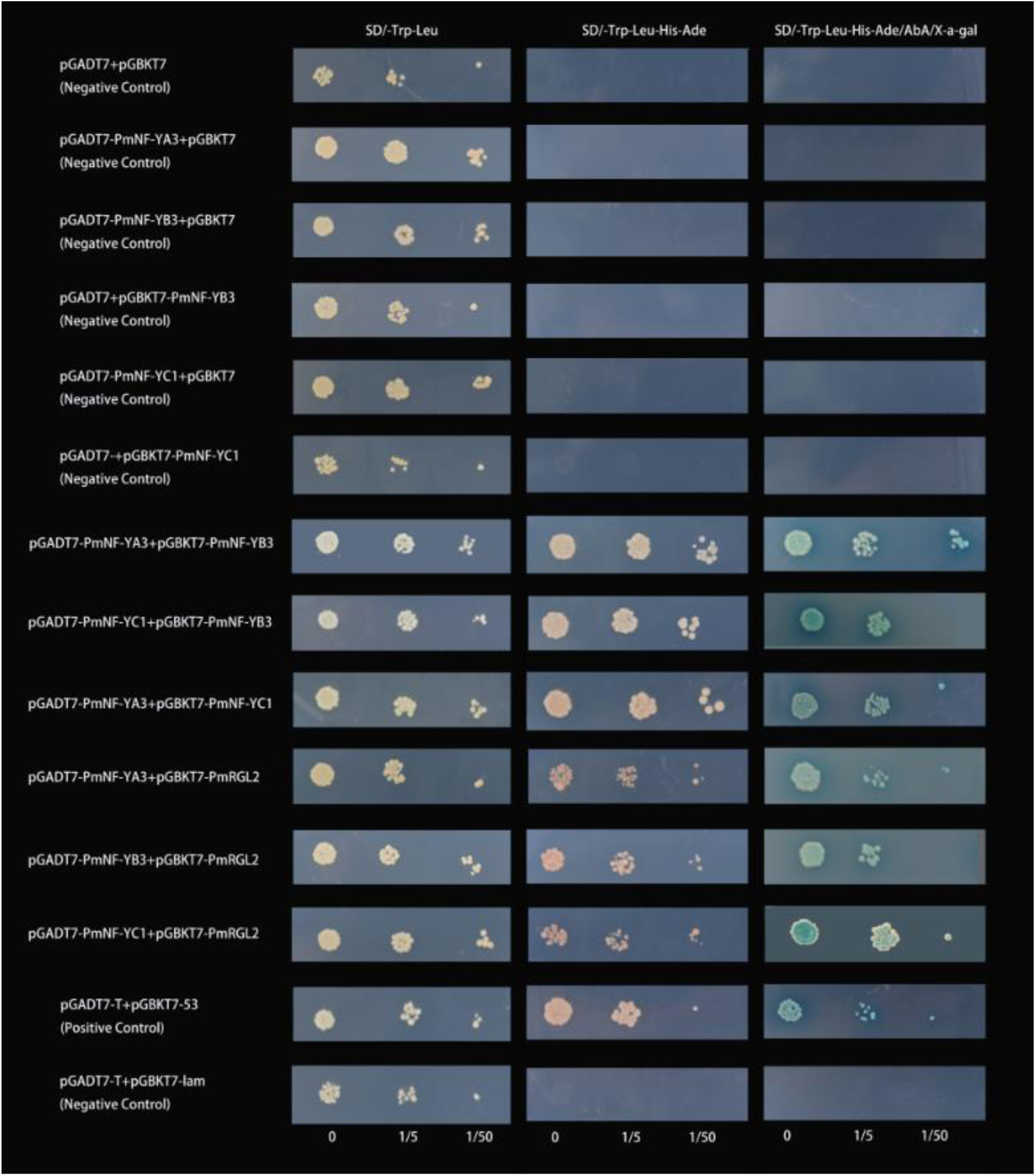
Interaction in yeast using the two-hybrid system. Notes: The pGBKT7 plasmid containing the CDs of the yeast transcription factor *NF-YA3, NF-YB3, NF-YC1* and *RGL2* (BD) was introduced into the Y2H Gold strain by transformation, whereas pGADT7, containing the CDs of *NF-YA3, NF-YB3, NF-YC1* and *RGL2* (AD), was introduced into the Y187 strain. These genes were mated in the combinations indicated and selected in synthetic defined media (SD) lacking Leu, Trp (SD-Trp-Leu, DDO) or Leu, Trp, adenine, and lacking His (SD-Trp-Leu-His-Ade, QDO) or lacking Leu, Trp, adenine, and His whit AbA (100ug/ml) and X-α-gal (QDO/A/X). Positive and negative controls are p53, interacting with AgT and LamC, respectively. Three dilutions (0, 1/5 and 1/50) of yeast cultures adjusted to an OD600 of 0.2 were spotted on synthetic medium to maintain proper growth. Photos were taken after four days of culture.

**Figure 9:**
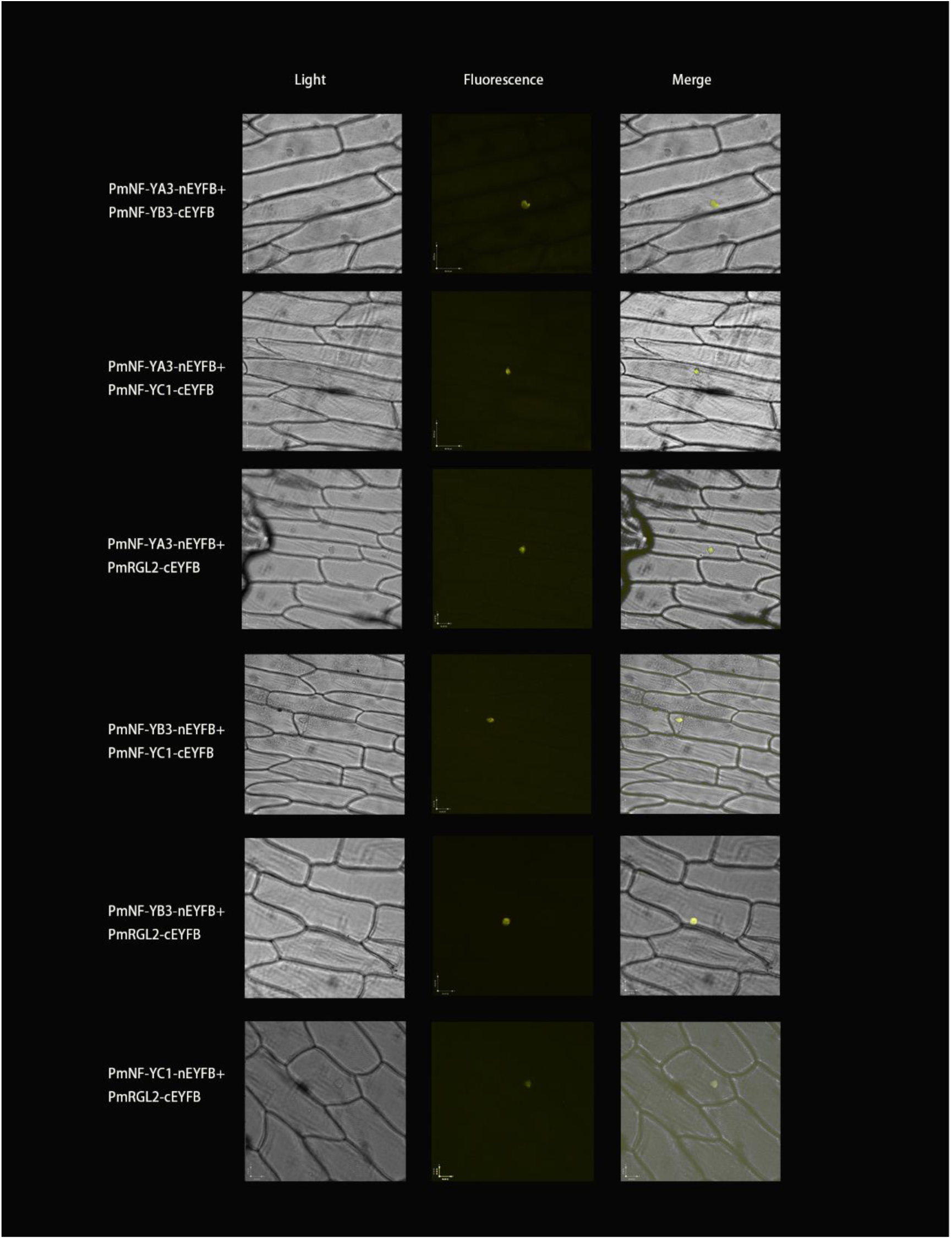
BiFC analysis of the interactions among NF-Y subunits and between NF-Y subunits and *PmRGL2* in onion epidermal cells. Notes: The same image, obtained with a confocal microscope, is shown under white light (left), UV (middle), and merged (right). Bar = 35μm.

## Discussion

### *PmRGL2* interacting with the NF-Y complex induces dormancy release in response to GA signals

Although it has been reported for a long time that GA, as a dormancy breaker, can replace part of the chilling requirements so that buds can break dormancy in advance (Saure, 1985). The mechanism of its action in dormancy is still unclear. DELLA proteins, as inhibitors of GA signaling associated with NF-Y in many reports. In tobacco, studies have shown that GA_3_ treatment increases the content of *AsNF-YB3*, which is also regulated by ABA and MeJA (Sun et al., 2016). In *Arabidopsis thaliana*, recent studies have shown that the NF-Y complex participates in GA-induced dormancy transition. REPRESSOR OF ga1-3(RGA) inhibits flowering by inhibiting NF-Y and CO interaction to regulate flowering (Xu et al., 2016). Liu et al. (2016) found that NF-YC and *RGL2* codependently inhibited GA-mediated seed germination. He stressed different combinations of DELLA-NF-Y(C) that function in different biological processes. However, so far, it has been rarely reported in the dormancy of woody plants. Combined with our previous reports we suspected NF-Y might also plays a role in *PmRGL2*-induced dormancy in Japanese apricot.

In our study, we treated Japanese apricot sticks with GA_4_ and found that the expression level of *PmRGL2* was significantly suppressed, while the expression level of the NF-Y family was significantly increased compared with the untreated group. Then we found that the NF-Y family can interact with GA signal suppressor *PmRGL2* by yeast hybrid experiment and BiFC assay. Previous studies have shown that a DELLA ubiquitin E3 ligase complex (SCF^SLY1/GID2^) which targets DELLA protein for degradation by the 26S proteasome, a proteasome stimulated by gibberellin (Fu et al., 2004; Ariizumi et al., 2011). Moreover, *PmSLY1* has been shown to target and degrade *PmRGL2* in Japanese apricot (Lv et al., 2018). GA_4_ treatment degrades *PmRGL2*, which enhances NF-Y binding to germination-related genes (Fig. 7). Here we show a NF-Y complex composed of three different types of subunits, *NF-YA3, NF-YB3*, and *NF-YC1*, which in response to GA signals through interacted with *PmRGL2* controls dormancy release in Japanese apricot.

Our finding of *PmNF-YC1* is a common partner of other NF-Y subunits and *PmRGL2*. Moreover, it has a similar expression pattern in each stage of dormancy with *PmRGL2*, which implies that *PmNF-YC1* could be a key subunit that coordinates the perception of GA_4_ signals and the assembly of the specific combinatorial NF-Y transcription factor that triggers the downstream regulatory events in response to GA signals (Fig. 6). Recent research has shown that NF-YC serves as a key component that not only mediates the interaction between NF-YA and NF-YB in the NF-Y heterotrimer but also interacts in vivo with RGA (Hou et al., 2014). Moreover, plant NF-YC enables translocation of NF-YB to the nucleus, and that the resulting NF-YB/NF-YC heterodimer is required for further recruitment of NF-YA in the final assembly of the heterotrimeric NF-Y complex (Hackenberg et al., 2012). These indicate that the formation of the *RGL2* complex with NF-Y is based on the binding of NF-YC.

So that revealsthat NF-Y transcriptional factors play a critical regulatory role in signal transduction and dormancy release. DELLA proteins, such as *RGL2*, affect the bud growth through preventing NF-Y binding to a germination-related gene (Fig. 10).

**Figure 10.**
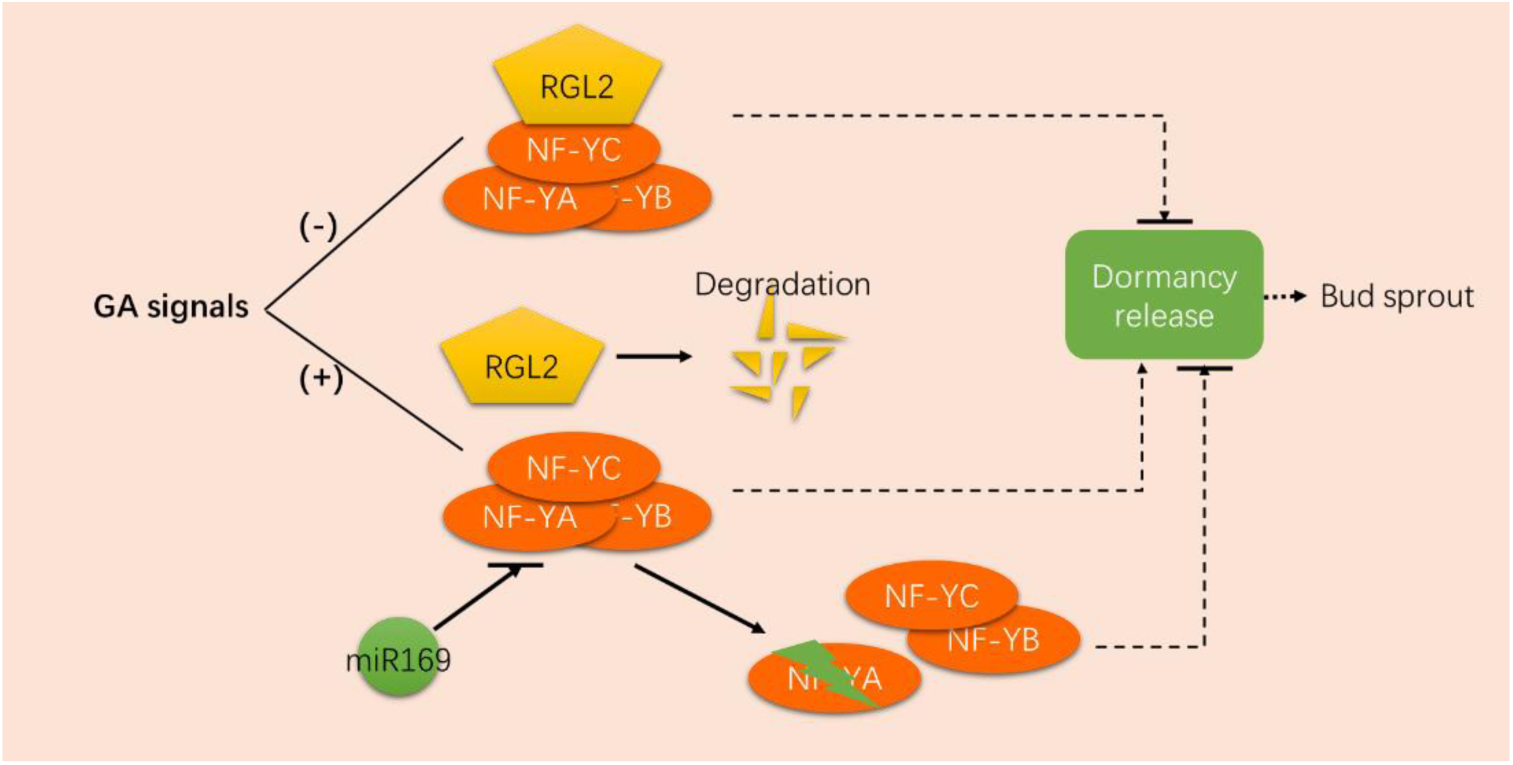
A working model of miR169 and *RGL2* synergism onthe NF-Y complex involved in the regulation of dormancy. Notes: In the absence of GA, *RGL2* binds to NF-Y to form as a complex inhibiting the release of dormancy in plants. When the GA signal is present, *RGL2* is degraded, and NF-Y complex targets downstream gene regulation dormancy. However, at low temperatures, a large number of miR169s are generated to target and degrade the NF-YA gene and destroy the function of the NF-Y complex.

### miR169 targets *PmNF-YA3* destruction of the NF-Y complex under the low temperature and inhibits the dormancy release

In general, a single miRNA can target multiple mRNAs and a single mRNA may be regulated by multiple miRNAs, this leads to a large number of potential regulatory effects (Kunz et al., 2014; Lee et al., 2018). MiR169 has been reported to play a role in multiple biological pathways in plants, and it works primarily by targeting NF-YA and cutting it (Hanna et al., 2010; Francesco et al., 2011; Gyula et al., 2018). MiR169 and NF-YA have been reported to play a crucial role in various life processes (Meng et al., 2011; Jean-Philippe et al., 2006; Céline et al., 2014; Mingda et al., 2015; Ni et al., 2013). Besides, Rewati et al.(2013) found that miR169 targets *PtrHAP2* and participates in the bud dormancy process of poplar. Ding et al.(2016) have shown that miR169 targeting *PagHAP2-6* is regulated by ABA in poplar dormancy. At present, there are few reports of miR169s in dormancy, and the mechanism of its participation in dormancy is still unknown.

Sequencing results showed that miR169s remained at a high-level during dormancy and were downregulated during dormancy release, which was consistent with RT-qPCR. Furthermore, it was predicted to target NF-YA through multiple software. Here we verified the methods of Dmitriev et al. (2013), which combined miRNA and mRNA sequencing results to verify the targeting relationship of miRNAs. Then we used 5’ RLM-RACE to demonstrate that Pmu-miR169e_3 targets *PmNF-YA3* in Japanese apricot as previously reported in other species, such assoybeans, poplar, Arabidopsis et al.(Ni et al., 2013; Ding et al., 2016; Du et al., 2017). We examined the expression of *PmNF-YA3* at different stages of dormancy. During dormancy period, its expression level decreased, which was in contrast to the miR169s expression levels. This is consistent with previous reports that miR169s may regulate the dormancy process by targeting and inhibiting *PmNF-YA3* (Céline et al., 2014). It was demonstrated that miR169s targeted and cleaved *PmNF-YA3* to inhibit its expression, thus inhibiting the release of dormancy in Japanese apricot.

Most temperate deciduous trees require a low-temperature exposure to be released from endodormancy. This is called chilling requirement (Fan et al., 2010; Kitamura et al., 2018). If the plant’s chilling requirement not sufficient, budburst will be delayed, reduced, or uneven budburst and flowering (Erez et al., 2000). When plantsare in the endodormancy period, dormancy can not be released even if the heat requirement sufficient. However, when the chilling requirement accumulated to a certain extent, GA_4_ treatment can effectively advance germination (Zhuang et al., 2013). This indicates that GA_4_ can replace part of the chilling requirement. Besides, the previous studies demonstrated that miR169s responded to environmental factors and further demonstrated that it could respond to low-temperature signals (Hanna et al., 2010; Gyula et al., 2018). These observations all support that at low temperatures, a large number of Pmu-miR169s are generated to target and degrade the *PmNF-YA3* gene and destroy the function of the NF-Y complex. When the heat requirement was sufficient, the amount of Pmu-miR169s will be reduced and the *PmNF-YA3* geneforms as NF-Y complex with two other subgroups to the regulation of dormancy. However, if the chilling requirement does not sufficient, even if adequate heat requirement, the NF-Y complex can not release dormancy due to the action of *PmRGL2*. However, when GA_4_ treatment replaces part of the chilling requirement, *PmRGL2* degrades, and heat requirement meet the demand, the NF-Y complex will act on the downstream gene and regulated the dormancy release. In other words, when the temperature is appropriate and GA is present, the *PmRGL2* degrade because of the generation of GA signals and MiR169s decreases because of the heat requirement, then the NF-Y complex can play a role in promoting dormancy release and promote plant bud sprout. The temperature and GA are both indispensable in dormancy (Fig. 10).

### *PmRGL2* and NF-Y complex may regulate ABA signaling as well as GA signaling in dormancy release

ABA is involved in different life processes as a plant hormone (Pei et al., 1998; Y, Ma et al., 2018; Ghate et al., 2019). It is generally considered to be an inhibitor of bud dormancy (Bris et al., 1999; Wen et al., 2016). ABA and GA antagonistically control seed dormancy (Yamaguchi et al., 2007; Chen et al., 2019). Zhang et al. (2018) used transcriptome to elucidate the relationship between hormone and sugar changes in different stages of dormancy and believed that GA and ABA were crucial in dormancy. However, the molecular mechanism of the relationship between ABA and GA remains unclear. Previous studies in our group determined the ABA, GA_3_ and GA_4_ content of flower buds in four stages of dormancy of Japanese Apricot (Wen et al., 2016). ABA concentrations significantly reduce in flower buds during induced dormancy stageand dormancy-release stage. The concentrations of GA_3_ and GA_4_ continue to rise until the dormancy release reaches its maximum. In transcriptome results, Pmu-mi166u increased during the dormancy period and decreased during the dormancy release period, which was predicted the targeting of the ABA-responsive element binding factor. The trend of miRNAs, which was predicted to target the GA related gene, was reversed to Pmu-mi166u (Supplementary Table S3).

Furthermore, the regulation of *RGL2* and NF-YCs is highly correlated and interdependent, forming complexes rather than binding to ABI5’s CCAAT domain alone to regulate *Arabidopsis* seed germination (Liu et al., 2016). It is suggested that the NF-Y complex may be involved in ABA signal transduction and act as a medium for ABA and GA signal connection. DELLA and NF-Y form a complex that binds to ABA signal transduction factor ABI5 to connect GA and ABA signaling pathways and thus play a role in dormancy transition. This will provide a new idea for understanding the mechanism of dormant molecules, in the absence of GA, *PmRGL2* binds to NF-Y to form as a complex and promotes related genes of ABA conduction, like *ABI5*, thus inhibiting the release of dormancy in plants.

## Conclusion

In this study, we found that Pmu-miR169s and its target gene *PmNF-YA3* contributed regulation of the dormancy release induced by gibberellin. GA_4_ could inhibit the expression of *PmRGL2* but promote the expression of the NF-Y gene during the dormancy release. Moreover, *PmRGL2* could interact with NF-Y family genes based on yeast two-hybrid and bimolecular fluorescence complementation assays. These results indicated that miR169 and *PmRGL2* synergistically regulate the NF-Y complex to active dormancy release in Japanese apricot, which provides the novel idea to discover the molecular mechanism of dormancy release in woody plants.

## MATERIAL AND METHODS

### Plant materials and sampling

The flower buds of Japanese Apricot cv. ‘Taoxingmei’, a low chilling requirement cultivar, and ‘Bungo’, a high chilling requirement cultivar, were collected from the National Field GenBank for Japanese apricot located in Nanjing, Jiangsu Province, during paradormancy, endodormancy, ecodormancy and dormancy release periods. The dormancy status of ‘Taoxingmei’ and ‘Bungo’ was investigated by our research group. The sampling dates were determined according to chilling hours and phenological traits (Gao et al. 2012b). The flower buds were collected from three trees, frozen in liquid nitrogen, and kept at -70°C until RNA isolation.

### RNA extraction, small RNA library construction, and sequencing

Samples from the flower buds at the four critical dormancy stages were pooled for RNA isolation and library construction. The total RNA isolated from the stages of paradormancy, endodormancy, ecodormancy, and dormancy release was named libraries Para, Endo, Eco, and Dr Each collected was performed with three biological replicates for high-throughput RNA-sequencing, respectively. RNA extraction was performed according to the manufacturer’s instructions using Trizol reagent (Invitrogen, USA). The concentration of RNA samples was analyzed using NanoDrop ND-1000 spectrophotometry (NanoDrop Technologies, Rockland, DE, USA) and Agilent 2200 TapeStation (Agilent Technologies, USA) in order to ensure the high quality of sample for sequencing. The amplified products were purified by gel electrophoresis and using BGISEQ-500 technology via the Illumina Genome Analyzer at the Beijing Genomics Institute (BGI, Shenzhen, China).

### Bioinformatics analysis

For members of the miR169 gene family from the miRBase (University of Manchester, http://www.mirbase.org/). Clean tags were mapped to the sRNA database such as miRBase, Rfam (http://rfam.xfam.org/), siRNA, piRNA, snoRNA, etc. Pmu-miR169 phylogenetic tree analysis and reliability values at each branch representing bootstrap samples (1000 replicates) were generated using MEGA5 software (Tamura et al., 2011). Prediction of mature miR169 family member target sequences was performed by psRobot, TAPIR\reference(TAPIR_ref), and TargetFinder. Combined with the corresponding filtering conditions such as free energy and score value, the results supported by at least two kinds of target gene prediction software were selected.

### 5’ RACE analysis

We use psRobot (http://omicslab.genetics.ac.cn/psRobot/), TAPIRor TargetFinder (https://github.com/carringtonlab/TargetFinder) to predict targets. The cleavage site in the miR169 gene was verified by 5′ RLM-RACE using a FirstChoice® RLM-RACE Kit (Ambion) according to the manufacturer’s instructions. The 5′ RACE cDNA was synthesized from mRNA ligated with a 5′ adaptor using random decamers. Nested PCR was performed to obtain the 3′ cleavage products of the target gene using a set of adaptors and gene-specific primers. The 5′ RACE cDNA was also used to detect the accumulation of 3′ cleavage products by polymerase chain reaction (PCR). Adaptors and adaptor primers were provided with the kit. Gene-specific primers are listed in Supplementary Table S1. All PCR products were cloned into the pMD19-T Easy vector (TAKARA) and sequenced.

### GA_4_ treatment assay

From December 20, 2018, annual shoots with full buds and 20-40 cm in length were collected in the field within seven days and quickly brought back to the laboratory wrapped in a wet cloth. The branches were divided into two groups and cultured in a light incubator (RXZ-1000B) with 200 μmol·L^-1^ GA_4_ and distilled water, respectively (with about 40 flower buds per branch and five branches as a plot. Repeat tentimes. With temperature (21±3°C), light 12h/ dark 12h, light intensity 55 μmol·m^-2^·s^-1^, air humidity 70%. When the flower bud germination rate on the branch reached 50%, the branch dormancy was considered to be lifted (Zhuang et al., 2013). The flower buds collected from the base of branches treated at 0, 3, 5, 8 and 10 days, freeze with liquid nitrogen and stored in the refrigerator at -70°C.

### Yeast two-hybrid assay

The full-length CDs of *PmNF-YA3, PmNF-YB3*, and *PmNF-YC1*, were amplified by PCR with gene-specific primers and inserted into pGADT7 and pGBKT7 vector, respectively (Supplementary Table S1). These constructs were transformed into Y2H Gold cells following the manufacturer’s protocol (Clontech). Self-activation and toxicity detection of the recombinant plasmid and control vectors (pGADT7-T, pGBKT7-53, and pGBKT7-Lam) were carried out as described in the Matchmaker TM Gold Y2H manual. Cultured yeast cells re-suspended in YPDA were plated on selective DDO media and incubated at 30°C for 3-5 days. Yeast cultures containing either the interactor vector, positive vector (pGADT7-T+pGBKT7-53), or negative vector (pGADT7-T+pGBKT7-Lam) were all selected in synthetic defined media (SD) lacking Leu, Trp (SD-Trp-Leu, DDO) or Leu, Trp, adenine, and lacking His (SD-Trp-Leu-His-Ade, QDO) or lacking Leu, Trp, adenine, and His whit AbA (100ug/ml) and X-α-gal (QDO/A/X). Positive and negative controls are p53, interacting with AgT and LamC, respectively. Three dilutions (0, 1/5 and 1/50) of yeast cultures adjusted to an OD600 of 0.2 were spotted on synthetic medium to maintain proper growth. These genes were mated in the combinations indicated and selected at 30°C for 3-5 days. Positive colonies were confirmed by PCR.

### Bimolecular fluorescence complementation assay

In the BiFC constructs, the full-length CDs of *PmNF-YA3, PmNF-YB3, PmNF-YC1*, and *PmRGL2* were inserted into pSAT4A-nEYFP-N1 and pSAT4A-cEYFP-N1 vector at the *SacII*/*BamHI* cloning sites using the primer pair, respectively. Transient gene expression in onion epidermal cells was performed using a Biolistic PDS-1000/He Particle Delivery System (Bio-Rad) according to the manufacturer’s instructions. After bombardment, the onion peels wereincubated at 25 °C for 16 h-18h in the dark. YFP fluorescence images were acquired by a 3D live cell imaging system (model Ultra VIEW VoX). Yellow fluorescence was photographed in light, fluorescence, and merge fields. All primers used in bimolecular fluorescence complementation assays are listed in Supplementary Table S1.

### Analysis of gene expression

Stem-loop real time quantitative PCR (RT-qPCR) was used to determine the relative levels of mature miR169 with 5.8 s rRNA as an internal control. The transcript levels of Pmu-miR169, the target gene, and other genes were quantified using RT-qPCR and normalized using RP II as a housekeeping gene. RT-qPCR was performed using the ABI 7300 Real-Time PCR System (Applied Biosystems, Foster City, CA, USA) and SYBR Green Real-time PCR Master Mix (Toyobo, Osaka, Japan). The experiments were performed using the method described above. Each analysis consisted of three biological replicates and three technical replicates of each biorep. The conditions for the PCR amplification were as follows: 95 °C for 3 min, followed by 40 cycles of 95°C for 20 s, 60°C for 20 s, and 72°C for 43 s. The relative expression level of genes and miRNAs wascalculated using the 2^-ΔΔCT^ method (Wu et al., 2017).

### Statistical analysis

The experiments were repeated three times, with three replicates for each. All data, expressed as mean ± standard errors (SE; n=3), were analyzed using SPSS 17.0 software (SPSS Inc., Chicago, IL). A comparison between groups was conducted by analysis of variance (ANOVA). Differences between test samples were determined using Duncan’s multiple range test at a significancelevel of P≤0.05 and extremely significant level of P≤0.01.

## Acknowledgment

The authors thanks to the National Key Research and Development Program of China (2018YFD1000107), the Priority Academic Program Development of Jiangsu Higher Education Institutions (PAPD) and the National Natural Science Foundation of China (31772282 and 31971703) for funding this research in materials collection, data analysis, and experiment.

## References

Acheampong AK, Chuanlin Z, Tamar H, Lisa G, Yumiko T, Yusuke J, Yuji K, Amnon L, Etti O (2017) Abnormal Endogenous Repression of GA Signaling in a Seedless Table Grape Cultivar with High Berry Growth Response to GA Application. Frontiers in Plant Science 8: https://doi.org/10.3389/fpls.2017.00850

Antonella R, Marianna B, Nicola M, Roberto M (1995) CCAAT-box binding protein NF-Y (CBF, CP1) recognizes the minor groove and distorts DNA. Nucleic Acids Research 23: 4565–4572.

Ariizumi, Tohru, Author, Lawrence, Paulraj, K., Author, Steber, Camille, M. (2011) The Role of Two F-Box Proteins, SLEEPY1 and SNEEZY, in Arabidopsis Gibberellin Signaling. Plant Physiology 155: 765–775.

Axtell MJ, Meyers BC (2018) Revisiting criteria for plant miRNA annotation in the era of big data. Plant Cell 30: 272–284.

Benmoussa HF, Ghrab M, Ben Mimoun M, Luedeling E (2017) Chilling and heat requirements for local and foreign almond (Prunus dulcis Mill.) cultivars in a warm Mediterranean location based on 30 years of phenology records. Agricultural & Forest Meteorology 239: 34–46

Brian PW, Hemming HG, Radley M (1955) A Physiological Comparison of Gibberellic Acid with Some Auxins. Physiologia Plantarum 8: 899–912

Bris ML, Michaux-Ferrière N, Jacob Y, Poupet A, Barthe P, Guigonis JM, Page-Degivry MTL (1999) Regulation of bud dormancy by manipulation of ABA in isolated buds of Rosa hybrida cultured in vitro. 26: 273–281

Campoy, J. A, Ruiz, EGEA (2011) Dormancy in temperate fruit trees in a global warming context: A review. Scientia Horticulturae 130: 357–372

Carnac G (2013) Simultaneous miRNA and mRNA transcriptome profiling of human myoblasts reveals a novel set of myogenic differentiation-associated miRNAs and their target genes. Bmc Genomics 14: 1–20 http://www.biomedcentral.com/1471-2164/14/265

Chen H, Ruan J, Chu P, Fu W, Huang S (2019) AtPER1 enhances primary seed dormancy and reduces seed germination by suppressing the ABA catabolism and GA biosynthesis in Arabidopsis seeds. The Plant Journal 101: 310–323

Chen H, Yang Q, Chen K, Zhao S, Zhuang W (2019) Integrated microRNA and transcriptome profiling reveals a miRNA-mediated regulatory network of embryo abortion under calcium deficiency in peanut (Arachis hypogaea L.). BMC Genomics 20: 1–17 https://doi.org/10.1186/s12864-019-5770-6

Chen L, Xiang S, Chen Y, Li D, Yu D (2017) Arabidopsis WRKY45 interacts with the DELLA protein RGL1 to positively regulate age-triggered leaf senescence. Molecular Plant 10: 1174–1189

Christophe R, Fabienne C, Roberto M, Dino M (2003) The NF-YB/NF-YC structure gives insight into DNA binding and transcription regulation by CCAAT factor NF-Y. Journal of Biological Chemistry 278: 1336–1345

Curaba J, Talbot M, Li Z, Helliwell C (2013) Over-expression of microRNA171 affects phase transitions and floral meristem determinancy in barley. Bmc Plant Biology 13: https://doi.org/10.1186/1471-2229-13-6

Ding Q, Zeng J, He X (2016) MiR169 and its target PagHAP2-6 regulated by ABA are involved in poplar cambium dormancy. Journal of Plant Physiology 198: 1–9

Dmitriev P, Barat A, Polesskaya A, O Connell MJ, Robert T, Dessen P, Walsh TA, Lazar V, Turki A, Du Q, Zhao M, Gao W, Sun S, Li WX (2017) microRNA/microRNA* complementarity is important for the regulation pattern of NFYA5 by miR169 under dehydration shock in Arabidopsis. Plant Journal 91: 22–33

Erez A (2000) Bud Dormancy; Phenomenon, Problems and Solutions in the Tropics and Subtropics. Temperate Fruit Crops in Warm Climates. In: Erez A (ed) Temperate fruit crops in warm climates. Kluwer, Dordrecht, pp 17–48

Fan S, Bielenberg DG, Zhebentyayeva TN, Reighard GL, Okie WR, Holland D, Abbott AG (2010) Mapping quantitative trait loci associated with chilling requirement, heat requirement and bloom date in peach (Prunus persica). New Phytologist 185: 917–930

Feng L, Daniela P, Claire B, Brunkard JO, Cohn MM, Jeffery T, Haoyu S, Pavan K, Barbara B (2012) MicroRNA regulation of plant innate immune receptors. Proceedings of the National Academy of Sciences of the United States of America 109: 1790–1795

Francesco L, Weits DA, Bikram Datt P, Wolf-Rüdiger S, Peter G, Dongen JTV (2011) Hypoxia responsive gene expression is mediated by various subsets of transcription factors and miRNAs that are determined by the actual oxygen availability. New Phytologist 190: 442–456

Fu X, Richards DE, Fleck B, Xie D, Burton N, Harberd NP (2004) The Arabidopsis Mutant sleepy1gar2-1 Protein Promotes Plant Growth by Increasing the Affinity of the SCFSLY1 E3 Ubiquitin Ligase for DELLA Protein SubstratesW. 16: 1406–1418

Gao Z, Shi T, Luo X, Zhang Z, Zhuang W, Wang L (2012a) High-throughput sequencing of small RNAs and analysis of differentially expressed microRNAs associated with pistil development in Japanese apricot. Bmc Genomics 13: https://doi.org/10.1186/1471-2164-13-371

Gao, Z, Zhuang, W, Wang, L, Shao, J, Luo, X, Cai, B, et al (2012b). Evaluation of chilling and heat requirements in Japanese apricot with three models. HortScience 12: 1826–1831

Ghate T, Barvkar V, Deshpande S, Bhargava S (2019) Role of ABA Signaling in Regulation of Stem Sugar Metabolism and Transport under Post-Flowering Drought Stress in Sweet Sorghum. Plant Molecular Biology Reporter 37: 303–313

Gyula P, Baksa I, Tóth T, Mohorianu I, Szittya G (2018) Ambient temperature regulates the expression of a small set of sRNAs influencing plant development through NF^TEL^A2 and YUC2. Plant Cell & Environment 41: 2404–2417

Hackenberg D, Wu Y, Voigt A, Adams R, Schramm P, GrimmI B (2012) Studies on Differential Nuclear Translocation Mechanism and Assembly of the Three Subunits of the Arabidopsis thaliana Transcription Factor NF-Y. Molecular Plant 5: 876–888

Hanna L, Seong Jeon Y, Jeong Hwan L, Wanhui K, Seung Kwan Y, Heather F, Carrington JC, Hoon AJ (2010) Genetic framework for flowering-time regulation by ambient temperature-responsive miRNAs in Arabidopsis. Nucleic Acids Research 38: 3081–3093

Hernandez Y, Sanan-Mishra N (2017) miRNA mediated regulation of NAC transcription factors in plant development and environment stress response. Plant Gene 11: 190–198

Horvath DP, Anderson JV, Chao WS, Foley ME (2003) Knowing when to grow: signals regulating bud dormancy. Trends in Plant Science 8: 534–540

Hou X, Zhou J, Liu C, Liu L, Shen L, Yu H (2014) Nuclear factor Y-mediated H3K27me3 demethylation of the SOC1 locus orchestrates flowering responses of Arabidopsis. Nature Communications 5: 1–14

Jagadeeswaran G, Sunkar ASAR (2009) Biotic and abiotic stress down-regulate miR398 expression in Arabidopsis. Planta 229: 1009–1014

Jean-Philippe C, Florian F, Fran Oise DB, Adnane B, Fikri EY, Sandra M, Tatiana V, Thomas O, Pascal G, Martin C (2006) MtHAP2-1 is a key transcriptional regulator of symbiotic nodule development regulated by microRNA169 in Medicago truncatula. Genes & Development 20: 3084–3088

Jiang N, Meng J, Cui J, Sun G, Luan Y (2018) Function identification of miR482b, a negative regulator during tomato resistance to Phytophthora infestans. Horticulture Research 5: https://doi.org/10.1038/s41438-018-0017-2

Jonesrhoades MW, Bartel DP, Bartel B (2006) MicroRNAs AND THEIR REGULATORY ROLES IN PLANTS. Annual Review of Plant Biology 57: 19–53

Jung JH, Lee S, Yun J, Lee M, Park CM (2014) The miR172 target TOE3 represses AGAMOUS expression during Arabidopsis floral patterning. Plant Science An International Journal of Experimental Plant Biology 215-216: 29–38

Kitamura Y, Habu T, Yamane H, Nishiyama S, Kajita K, Sobue T, Kawai T, Numaguchi K, Nakazaki T, Kitajima A (2018) Identification of QTLs controlling chilling and heat requirements for dormancy release and bud break in Japanese apricot (Prunus mume). Tree Genetics & Genomes 14: https://doi.org/10.1007/s11295-018-1243-3

Kunz M, Xiao K, Liang C, Viereck J, Pachel C, Frantz S, Thum T, Dandekar T (2014) Bioinformatics of cardiovascular miRNA biology. Journal of Molecular & Cellular Cardiology: S0022282814003873. https://doi.org/10.1016/j.yjmcc.2014.11.027

Lang G A. (1987) Endo-,para-,and ecodormancy:physiological terminologyand classification for dormancy reseach. Hort Sci, 22: 371–377.

Lee JY, Yun SJ, Jeong P, Piao XM, Kim W (2018) Identification of differentially expressed miRNAs and miRNA-targeted genes in bladder cancer. Oncotarget 9: 27656–27666

Liu X, Hu P, Huang M, Tang Y, Li Y, Li L, Hou X (2016) The NF-YC-RGL2 module integrates GA and ABA signalling to regulate seed germination in Arabidopsis. Nature Communications 7: 1–14

Luan M, Xu M, Lu Y, Zhang L, Fan Y, Wang L (2015) Expression of zma-miR169 miRNAs and their target ZmNF-YA genes in response to abiotic stress in maize leaves. Gene 555: 178–185

Lv L, Huo X, Wen L, Gao Z, Khalilurrehman M (2018) Isolation and Role ofPmRGL2in GA-mediated Floral Bud Dormancy Release in Japanese Apricot (*Prunus mume* Siebold et Zucc.). Frontiers in Plant Science 9: https://doi.org/10.3389/fpls.2018.00027

Ma L, Zhou L, Quan S, Xu H, Niu J (2019) Integrated analysis of mRNA-seq and miRNA-seq in calyx abscission zone of Korla fragrant pear involved in calyx persistence. BMC Plant Biology 19 https://doi.org/10.1186/s12870-019-1792-0

Mantovani R The molecular biology of the CCAAT-binding factor NF-Y. Gene 239: 15–27

Martyn GE, Quinlan KGR, Crossley M (2016) The regulation of human globin promoters by CCAAT box elements and the recruitment of NF-Y. Biochimica et Biophysica Acta 1860: 525–536

Meng Z, Hong D, Jian-Kang Z, Fusuo Z, Wen-Xue L (2011) Involvement of miR169 in the nitrogen-starvation responses in Arabidopsis. New Phytologist 190: 906–915

Ni Z, Hu Z, Jiang Q, Zhang H (2013) GmNFYA3, a target gene of miR169, is a positive regulator of plant tolerance to drought stress. Plant Molecular Biology 82: 113–129

Pei ZM, Ghassemian M, Kwak CM, Mccourt PM, Schroeder JI (1998) Role of Farnesyltransferase in ABA Regulation of Guard Cell Anion Channels and Plant Water Loss. Science 282: 287–290

Peng J, Carol P, Richards DE, King KE, Cowling RJ, Murphy GP, Harberd NP (1997) The Arabidopsis GAI gene defines a signaling pathway that negatively regulates gibberellin responses? Genes & Development 11: 3194–3205

Posner D, Pavich S. (2018) Methods for improving bud break. 99: 6174–6180.

Prudencio AS, Martínez-Gómez P, Dicenta F (2018) Evaluation of breaking dormancy, flowering and productivity of extra-late and ultra-late flowering almond cultivars during cold and warm seasons in South-East of Spain. Scientia Horticulturae 235: 39–46

Ravindran P, Verma V, Stamm P, Kumar PP (2017) A Novel RGL2-DOF6 Complex Contributes to Primary Seed Dormancy in Arabidopsis thaliana by Regulating a GATA Transcription Factor. Molecular plant 10: 1307–1320

Rewati P, Jill R, Victor B (2013) ptr-MIR169 is a posttranscriptional repressor of PtrHAP2 during vegetative bud dormancy period of aspen (Populus tremuloides) trees. Biochem Biophys Res Commun 431: 512–518

Rinne PILH, Annikki W, Jorma V, Linda R, Raili R, Jaakko KR, Christiaan VDS (2011) Chilling of dormant buds hyperinduces FLOWERING LOCUS T and recruits GA-inducible 1,3-beta-glucanases to reopen signal conduits and release dormancy in Populus. Plant Cell 23: 130–146

S., Nimisha, D., Kherwar, M. K, Ajay, B., Singh, K. Usha (2013) Molecular breeding to improve guava (Psidium guajava L.): Current status and future prospective. Scientia Horticulturae 164: 578–588

Salanenka Y, Verstraeten I, Löfke C, Tabata K, Friml J (2018) Gibberellin DELLA signaling targets the retromer complex to redirect protein trafficking to the plasma membrane. Proc Natl Acad Sci U S A 115: 1–6 https://doi.org/10.1073/pnas.1721760115

Saure MC (2011) Dormancy Release in Deciduous Fruit Trees. Horticultural Reviews 7: 239–300

Singh I, Smita S, Mishra DC, Kumar S, Singh BK, Rai A (2017) Abiotic Stress Responsive miRNA-Target Network and Related Markers (SNP, SSR) inBrassica juncea. Frontiers in Plant Science 8: https://doi.org/10.3389/fpls.2017.01943

Steven F, Clay HA, Katherine D, Finch-Savage WE (2014) Environment sensing in spring-dispersed seeds of a winter annual Arabidopsis influences the regulation of dormancy to align germination potential with seasonal changes. New Phytologist 202: 929–939

Sun X, Ren Y, Zhang X, Lian H, Zhou S, Liu S (2016) Overexpression of a garlic nuclear factor Y (NF-Y) B gene, AsNF-YB3, affects seed germination and plant growth in transgenic tobacco. Plant Cell Tissue & Organ Culture 127: 513–523

Suzuki H, Park SH, Okubo K, Kitamura J, Nakajima M (2009) Differential expression and affinities of Arabidopsis gibberellin receptors can explain variation in phenotypes of multiple knock-out mutants. Plant Journal 60: 48–55

Sven Eriksson HBTM (2006) GA4 Is the Active Gibberellin in the Regulation of LEAFY Transcription and Arabidopsis Floral Initiation. Plant Cell 18: 2172–2181

Tao W, Huitang P, Jia W, Weiru Y, Tangren C, Qixiang Z (2014) Identification and profiling of novel and conserved microRNAs during the flower opening process in Prunus mume via deep sequencing. Molecular Genetics & Genomics 289: 169–183

Wen LH, Zhong WJ, Huo XM, Zhuang WB, Gao ZH (2016) Expression analysis of ABA- and GA-related genes during four stages of bud dormancy in Japanese apricot (Prunus mume Sieb. et Zucc). Journal of Pomology & Horticultural Science 91: 1–8

Wisniewski M, Fuchigami LH, Sauter JJ, Shirazi A, Zhen LP, Lang GA (1996) Near-lethal stress and bud dormancy in woody plants. In Plant Dormancy: Physiology, Biochemistry & Molecular Biology, 201–210

Wu X, Gong Q, Ni X, Zhou Y, Gao Z (2017) UFGT: The Key Enzyme Associated with the Petals Variegation in Japanese Apricot. Frontiers in Plant Science 8: https://doi.org/10.3389/fpls.2017.00108

Xu F, Li T, Xu PB, Li L, Du SS, Lian HL, Yang HQ (2016) DELLA proteins physically interact with CONSTANS to regulate flowering under long days in Arabidopsis. Febs Letters 590: 541–549

Y L, Y Y, Y H, H L, M H, Z Y, F K, X L, X H (2019) DELLA and EDS1 Form A Feedback Regulatory Module to Fine-tune Plant Growth-Defense Tradeoff in Arabidopsis. Molecular plant 12: 1485–1498

Y M, J C, J H, Q C, X L, Y Y (2018) Molecular Mechanism for the Regulation of ABA Homeostasis During Plant Development and Stress Responses. International journal of molecular sciences 19 https://doi.org/10.3390/ijms19113643

Yamaguchi S, Kamiya Y, Nambara E (2007) Regulation of ABA and GA Levels During Seed Development and Germination in Arabidopsis. In: Seed Development, Dormancy and Germination. Bradford KJ, Nonogaki H, eds., Blackwell, Oxford, pp. 224–247.

Yu X, Hou Y, Chen W, Wang S, Wang P, Qu S (2017) Malus hupehensis miR168 Targets to ARGONAUTE1 and Contributes to the Resistance against Botryosphaeria dothidea Infection by Altering Defense Responses. Plant & Cell Physiology 58: 1541–1557.

Zhan X, Wang B, Li H, Liu R, Kalia RK, Zhu JK, Chinnusamy V (2012) Arabidopsis proline-rich protein important for development and abiotic stress tolerance is involved in microRNA biogenesis. Proceedings of the National Academy of Sciences of the United States of America 109: 18198–18203

Zhang L, Chen L, Yu D (2018) Transcription Factor WRKY75 Interacts with DELLA Proteins to Affect Flowering. Plant Physiology 176: 790–803

Zhang X, Zhe Z, Gong P, Zhang J, Ziaf K, Li H, Xiao F, Ye Z (2011) Over-expression of microRNA169 confers enhanced drought tolerance to tomato. Biotechnology Letters 33: 403–409

Zhang Z, Zhuo X, Zhao K, Zheng T, Han Y, Yuan C, Zhang Q (2018) Transcriptome Profiles Reveal the Crucial Roles of Hormone and Sugar in the Bud Dormancy of Prunus mume. Scientific Reports 8: https://doi.org/10.1038/s41598-018-23108-9

Zhao B, Ge L, Liang R, Wei L, Jin Y (2009) Members of miR-169 family are induced by high salinity and transiently inhibit the NF-YA transcription factor. BMC Molecular Biology 10: https://doi.org/10.1186/1471-2199-10-29

Zheng C, Zhao L, Wang Y, Shen J, Zhang Y, Jia S, Li Y, Ding Z (2015) Integrated RNA-Seq and sRNA-Seq Analysis Identifies Chilling and Freezing Responsive Key Molecular Players and Pathways in Tea Plant (Camellia sinensis). Plos One 10: 1–28

Zhuang W, Cai B, Gao Z, Zhen Z (2016) Determination of chilling and heat requirements of 69 Japanese apricot cultivars. European Journal of Agronomy 74: 68–74

Zhuang W, Gao Z, Wang L, Zhong W, Ni Z, Zhang Z (2013) Comparative proteomic and transcriptomic approaches to address the active role of GA4 in Japanese apricot flower bud dormancy release. Journal of Experimental Botany 64: 4953–4966

